# Good wildfires in western U.S. forests (2010-2020)

**DOI:** 10.1101/2024.12.06.627082

**Authors:** Jennifer K. Balch, Tyler L. McIntosh, Nayani T. Illangakoon, Karen Cummins, J. Morgan Varner, Winslow D. Hansen, Kate Dargan Marquis, Crystal L. Raymond, Brian J. Harvey, Matthew E. Bitters, Rutherford V. Platt

## Abstract

Western U.S. forests have evolved with fire. However, recent wildfire disasters have understandably focused on the tragic consequences in the moment, lives lost and homes destroyed, rather than the longer-term ecosystem benefits. Some of the positive outcomes, particularly from low-moderate severity wildfires, are to reduce fuels and promote regeneration, even in the context of warmer temperatures (1). Here, we estimated the amount of “good wildfire” that burns in an ecologically beneficial way in forests, with a severity and return interval analogous to the historical fire regime. Of western U.S. forest wildfires (2010-2020), 58% burned as good wildfire (49% and 9% burned matching the low-moderate and high severity regimes, respectively). The low-moderate severity good wildfires burned 3.1 million forest ha (N = 18,061 events), more than double the 1.4 million ha of prescribed burning (N = 24,817 events on federal land) over the same period. At the event level, 56% of wildfires (N = 3479;>10% forest cover) had over half of their area burned as good wildfire—highlighting the opportunity that each fire presents for suppression teams to cultivate more beneficial wildfire in western landscapes. Good wildfires are an underrecognized but important tool to mitigate the risk of future wildfire disasters. With fires likely to increase in frequency and area with warming (2), the coming decade provides an essential opportunity to promote good wildfire for ecosystems, people, and property.

## Introduction

During wildfires, we understandably focus on the tragic consequences in the moment, lives lost and homes destroyed, rather than potential longer-term beneficial ecosystem outcomes. While wildfires can be disastrous for society, they can also restore forest structure and composition, improve habitat for imperiled biota, provide critical nutrients to soils, and reduce the extent and intensity of future fires that put communities and people at risk (2, 3). In effect, wildfires can be simultaneously detrimental and beneficial (4, 5). This dual nature of fire as a key ecological process and a threat to human communities forces us to reckon with how we appreciate fire in contemporary landscapes. This challenge is especially acute across western U.S. forest species that evolved with fire and often require it to maintain healthy structure and function (6–8). With the increasing trend in the size (9), higher severity area (10), and spread rates (11) of wildfires across the western U.S. in the past couple decades, particularly in our forest systems (12), we need to also acknowledge the positive aspects of burning.

Here, we quantified the amount of “good wildfire,” or ecologically beneficial wildfire, that has occurred across western U.S. forests (2010-2020). We define “good wildfire” as wildfires that burn with a severity and frequency that matches their known historical fire regime. We assume that wildfires burning within these parameters will be ecologically beneficial to the extant ecosystem. Extensive prior work shows how burning outside of historical frequency and severity has negative outcomes for forest structure, composition, and recovery (e.g., (13–15)), with strong evidence of regeneration after appropriate severity fires across the western U.S. (e.g., (16, 17, 1, 18)). To scale westwide, we developed a remote-sensing based approach (Figure S1) that leverages pixel-level observations (30 × 30 m) of both severity and return interval over the available satellite record (∼40 years) and then estimated the amount of forest area that has burned during wildfires with a severity and return interval that matches its historical fire regime. We consider this amount of area good wildfire, while wildfires burning at a severity or frequency outside their historical range are considered less likely to promote ecosystem recovery. In this approach, we used a conservative estimate of forest extent in the year prior to fire based on multiple land cover datasets, and measured contemporary fire severity using Landsat-derived estimates of the commonly-used Composite Burn Index (CBI) (see supplemental Methods) (19). We compared contemporary fire occurrence characteristics to severity and return interval historical ranges of variation as documented in the Landfire product suite, which represents the best available expert-derived wall-to-wall estimates of historical fire regimes. We further estimated the post-wildfire aboveground biomass density recovery of these selected fires using the aboveground biomass density (AGBD) metric derived from the Global Ecosystems Dynamics Investigation (GEDI) product, an explicit metric that can be used to investigate post-fire biomass change over a chronosequence matched to this study.

We focus here on ecologically beneficial wildfires that were reported, not intentional cultural burning that is part of Indigenous fire stewardship (20), termed ‘good fire’ by prominent Indigenous scholars and practiced by Tribal traditional knowledge holders for millennia (21, 22). In our focus on good wildfire, we delineate two classes: low-moderate severity wildfire in regions that historically burned this way, hereafter “lower severity regions,” and high severity wildfire in regions that historically had stand-replacing fires, hereafter “high severity regions.” Further, we compare this lower severity area of good wildfire with the amount of prescribed burning that has been conducted on forested public lands during the same decade. This comparison helps illustrate that wildfires are helping to reach treatment targets (and broader ecological and fire hazard reduction objectives), but are a largely unaccounted source of ecologically beneficial burning. While we focus on severity and return interval here, ecological benefits may be compromised when fire seasonality, size of high severity patches, the total size of a fire, and post-fire climate conditions diverge from historical norms (23, 1). Also, ecological benefits may be conferred by certain amounts of lower severity fire in regions with historically higher severity fire regimes and vice versa. Fires in these mixed severity regimes are dynamic and complex, contributing to high landscape-level diversity and providing habitat for species requiring different successional stages (24). The potential environmental benefits and interactions of both high- and low-severity fires in such mixed regimes are not fully accounted for here; as such, our results are likely underestimates of good wildfire. Additional data sources will offer opportunities to characterize these other dimensions of fire effects in the future (e.g., (25)). As fire management tactics shift from a focus on full suppression to more intentional forest restoration and proactive fuel treatments, we urgently need a recounting of wildfire to acknowledge the potential ecological benefits across western U.S. forests.

### More than half of wildfire burned area was aligned with historical severity and frequency (2010-2020)

We estimated that 58% of the forest area burned by wildfires from 2010 to 2020 was ecologically beneficial, or good wildfire. In effect, wildfires contributed a substantial, but unaccounted for, ecosystem benefit in that they burned analogously to their past fire regimes. Further, this amount was 2.7 times the forest area impacted by prescribed burning—supporting the contention that certain wildfires, or portions of them, should be counted in land management agencies’ restoration targets. The total area of lower severity good wildfire over this decade was 3.1 million ha for wildfires (N = 18,061 events; 49% of total forest area burned by wildfires), compared with 1.4 million ha for prescribed forest burns (N = 24,817 events) (Figure 1). Additionally, across the west (2010-2020), there were 589,000 ha of high severity good wildfire, representing an additional 9% of the total forest area burned during the time frame. Across all severities of good wildfire, this amounts to 3.7 million ha in the west (2010-2020). Only 3.4% of all forest wildfires that occurred from 2010-2020 burned more frequently than the estimated historical regime return interval. Among those fires, 32% burned too frequently as high severity (4 or 5H regimes) and 68% burned too frequently as low severity (3 or 5L regimes). On average, wildfires did beneficial work in 0.3 million ha per year in lower severity regions, but there was substantial interannual variation (annual minimum = ∼50,000 ha; annual maximum = ∼600,000 ha). This interannual pattern is largely due to differences in drought and non-drought years and the climatic controls on burned area (Figure 2), which is well documented (26). Only in extremely low fire years did low-moderate severity burning within wildfire perimeters constitute less area than that which occurred with prescribed burning.

**Figure 1.**
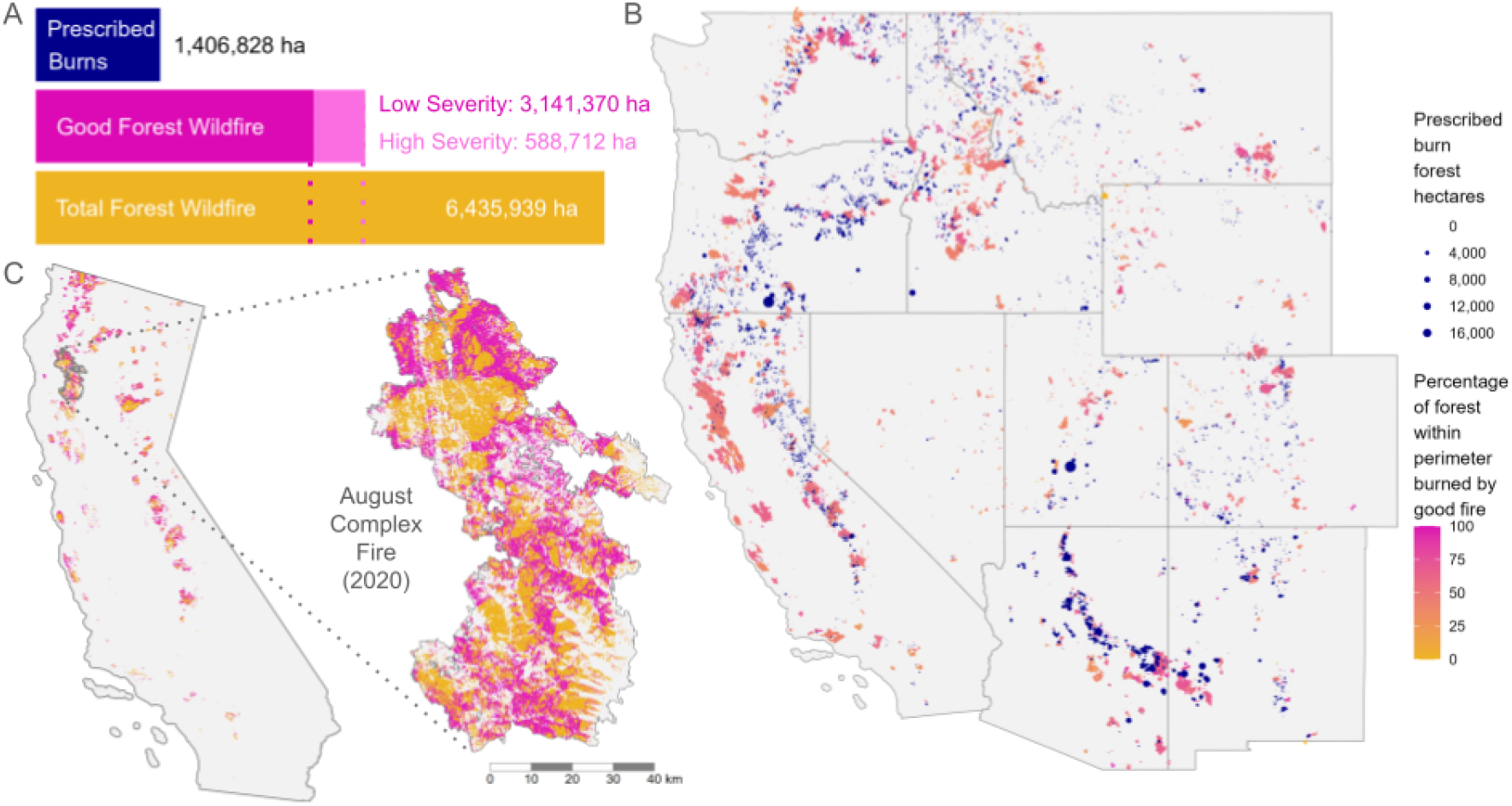
Good wildfire in western U.S. forests (2010-2020) **(A)** Area of prescribed fire, low-moderate and high severity good wildfire, and total wildfire. Good wildfire is a subset of total wildfire; prescribed fire is not. **(B)** The extent of good wildfire and proportion within individual fire perimeters (gradation from yellow (0%) to pink (100%)), compared with prescribed forest fires (blue). Fire perimeters contained >10% forest in the year before burn. **(C)** The extent of good wildfire in California and the August Complex fire showing the extent and patterning of good wildfire (pink) and all other forest fire (yellow).

**Figure 2.**
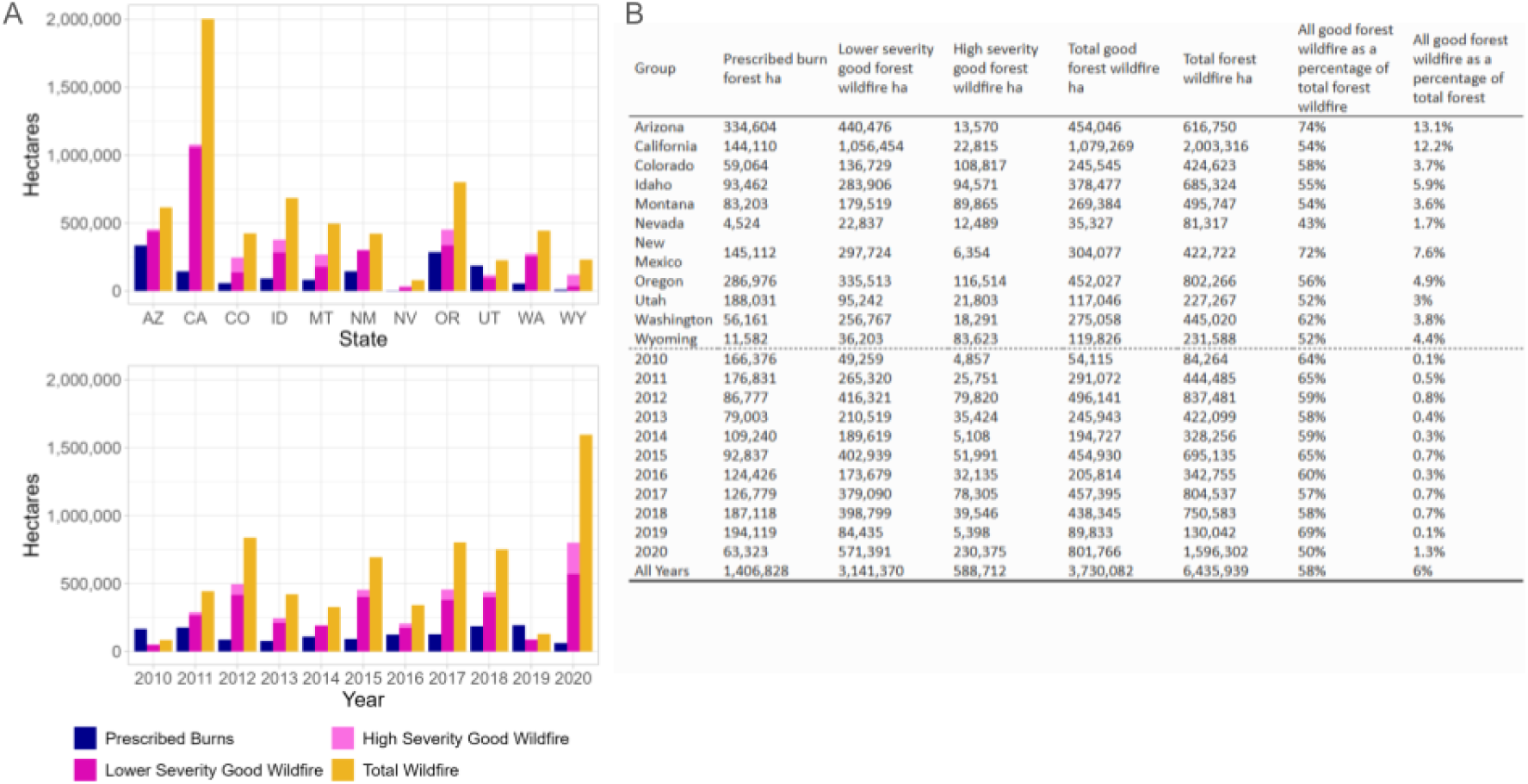
**(A)** The area (ha) of good wildfire in the western U.S. by state and year, compared with the total forest area that burned and the amount of prescribed fire conducted. Good wildfire is a subset of total wildfire; prescribed fire is not. **(B)** Area (ha) of total burned areas and good forest wildfire as a percentage of total forest wildfire and as a percentage of total forest. Total forest area for each state and ‘All Years’ is averaged over 2010-2020.

While there were substantial differences across states, the area of lower severity good wildfire was typically much higher than prescribed burning (Figure 2). California, which has experienced record-breaking wildfire years over the past decade, had over one million ha of lower severity good wildfire during this period, which was 7.3 times greater than what was accomplished through prescribed burning. Good wildfire is distributed in patches across individual fire events (e.g., the million acre (>400,000 ha) August Complex wildfire burned in a patchwork of low, moderate, and high severities, Figure 1C). That is, all wildfires are a mix of severities with some proportion that matches the historical fire regime. We performed whole fire-level analyses for events with >10% forested area to avoid spurious conclusions driven by fires that burned little forest. Fires with >10% forest make up 34% of all events analyzed, but account for 98-99% of all forested area burned and good wildfire area, respectively. In general, larger events had more intermediate percentages of good wildfire, while smaller events tended to have greater percentages of good wildfire (Figure 3A), reflecting the heterogeneity of both fire severity and fire regimes across scales. The median event had 54% good wildfire (Figure 3B). Event-level good wildfire distributions also varied by state: the median event in New Mexico had 68% good wildfire (highest), while the median value in Wyoming was 43% (lowest) (Figure 3C).

**Figure 3.**
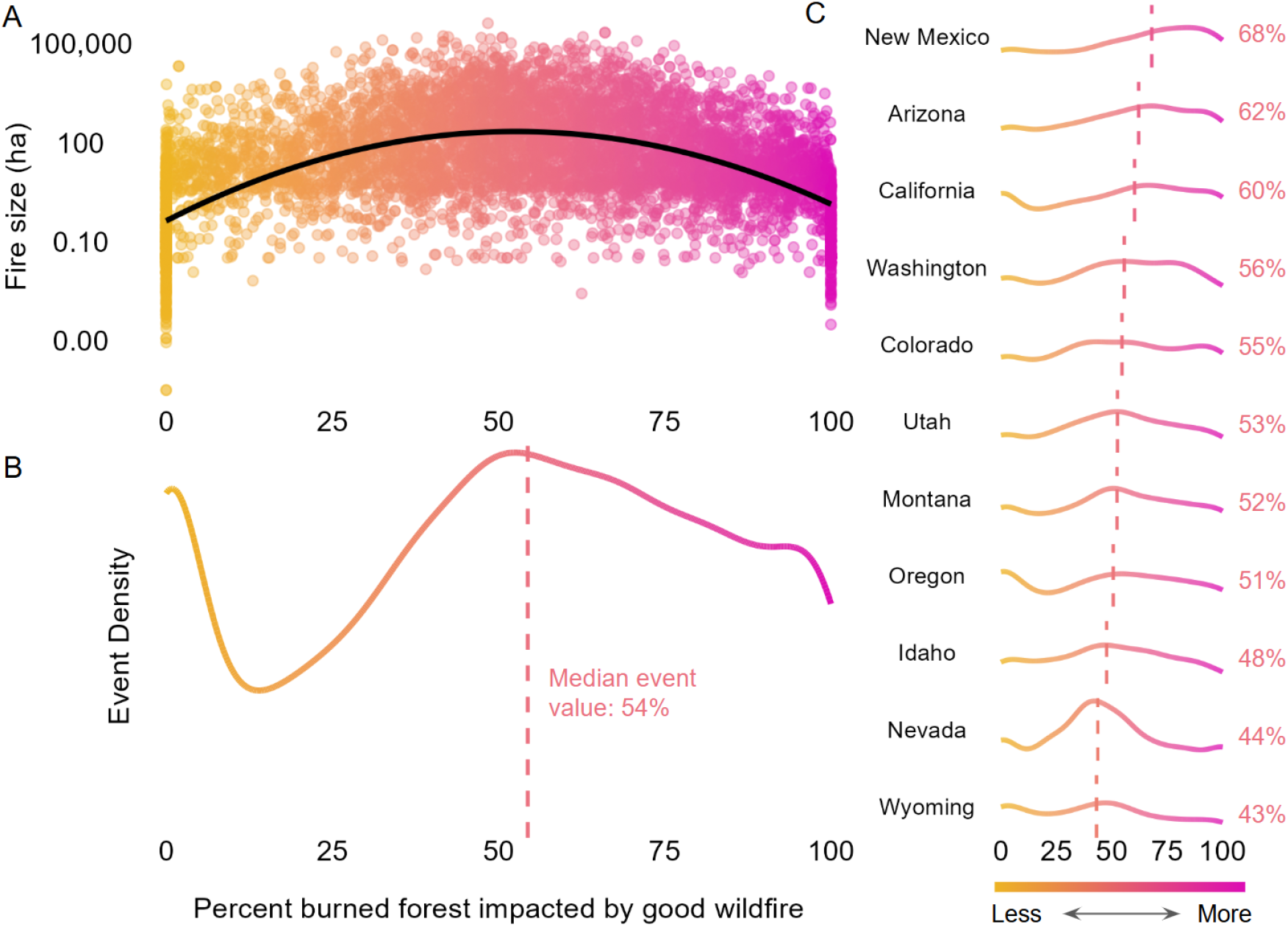
Percent good forest wildfire over events. All x-axes represent percent of burned forest impacted by good wildfire, and all panels show events with >10% forest in the year prior to burn. The orange to pink color scale represents the same x-axis in a more visual format (see color bar). **(A)** Scatter plot of fire size (log-transformed) against event-level percentage of good wildfire. Black line indicates a second order polynomial fit. **(B)** Density curve of percent good wildfire per event. Overall median value indicated by dashed pink line and annotation. **(C)** Density curves of percent good wildfire per event, subset by state. Median values indicated by dashed pink lines and annotations.

Of the forested prescribed burns included in our west-wide analysis, 77% by area occurred in lower severity regions. Sixty-four percent was designated as broadcast burns, i.e., intentional, controlled fire that burns through a predetermined landscape, while the remaining 36% was jackpot and pile burning (i.e., prescribed burning of fuels in scattered concentrations, and prescribed burning of hand- and machine-piled material, respectively) (27, 28). The area of prescribed burning reported here is a conservative estimate that only accounts for activity on federal lands and does not include state or private lands.

Our aboveground biomass density analysis showed that low severity fires burning in ways consistent with their historical fire regime had a lower percent change (<10%) in biomass and a faster recovery (∼5 years) towards baseline conditions (Figure 4).This reflects the potential for recovery faster than the historical fire return interval (≤35 years). Conversely, fires that burned outside their historical fire regime showed greater declines in biomass and slower recovery.

**Figure 4.**
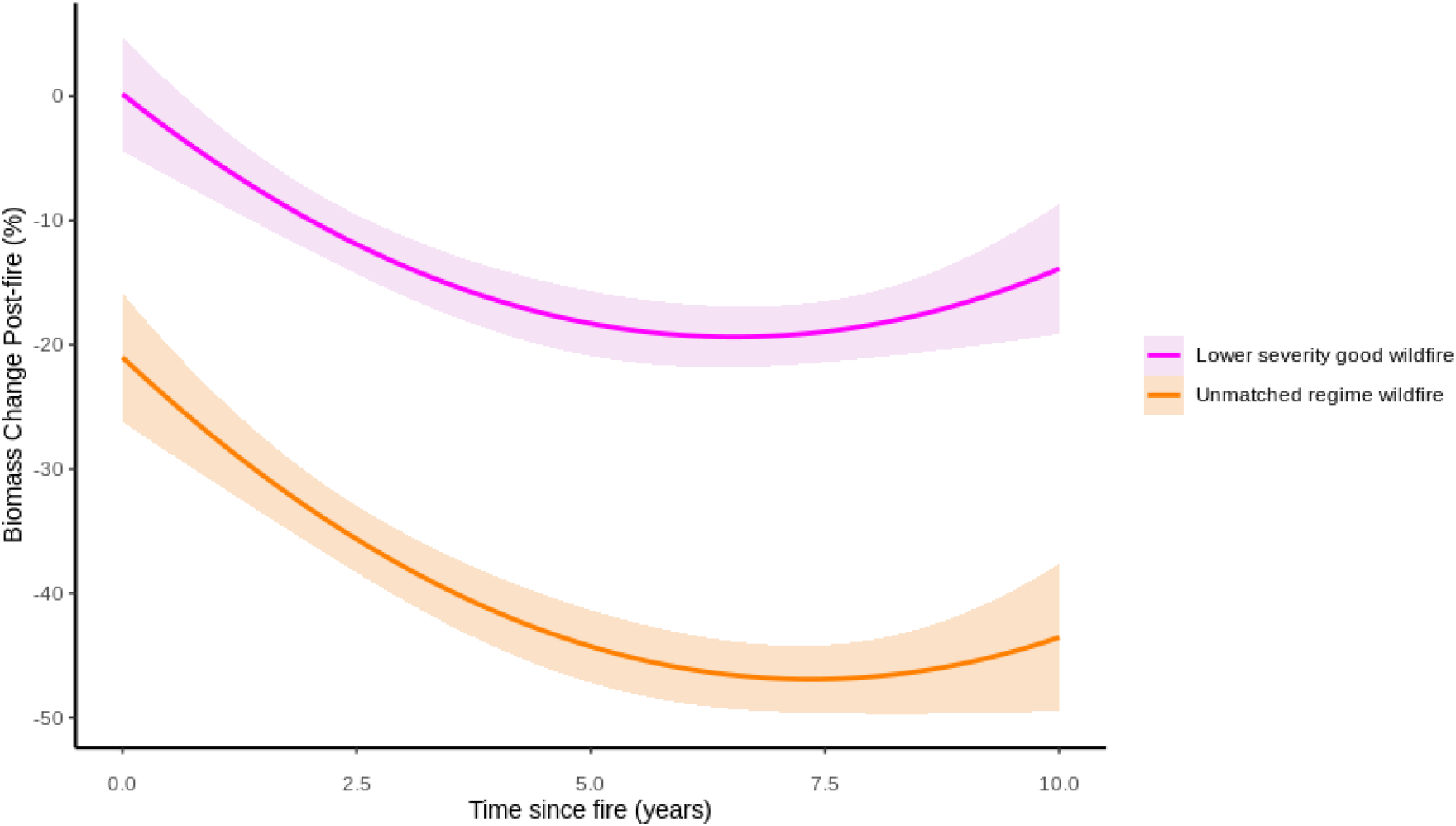
Percent biomass density change of burned areas with respect to unburned neighborhoods with time since fire. The solid lines represent the median and the shaded areas represent the standard deviation of biomass density. Pink indicates low and moderate severity fires that burned within its fire regime group. Orange indicates fires that burned with severity outside its fire regime group severity.

### Accounting for good wildfire is critical to stewarding burning back into our fire-adapted forests

This research recounts modern wildfires in the U.S. to emphasize their potential ecosystem benefits. Overall, our results demonstrate that 58% of the forest area burned by wildfires from 2010 to 2020 was potentially ecologically beneficial, or good wildfire. Moreover, this area was 2.7 times more hectares burned than accomplished through prescribed burning, demonstrating how substantial a contribution wildfires can make to treatment targets, such as the 2022 Wildfire Crisis strategy, which calls for two million hectares per year treated in a decade. Here, we focus only on wildfires that were reported and occurred with a severity and frequency analogous to the forests’ historical regime. By first accounting for good wildfire, we have the ability to understand when and where these types of wildfires occur and ultimately how land managers and fire response teams can better promote such fire in the future.

From a management perspective, fires that burned with high percentages of good wildfire may represent theoretical past opportunities for expanding the footprint of good wildfire on the landscape. That is, fires with moderately high percentages of good wildfire may have the potential for management to shift their dynamics towards those that are aligned with desired outcomes. Given that 98% of wildfires are suppressed before they become large, there is significant unrealized potential for achieving this (29). While the analysis presented here does not account for impacts on society and infrastructure, a high percentage of good wildfire in a single event indicates that fire behavior was likely well-aligned with desired outcomes. From 2010-2020, 56% of the wildfires occurring in areas with >10% forest cover (N = 3,479 / 6,257) had over half of their area burned as good wildfire, primarily in the form of lower severity good wildfire (85% of the good wildfire area in those events). Events with over half of their area as good wildfire events cumulatively make up 68% of the good wildfire area in forested events (Figure 5A). California had the highest number of these events, but they make up the highest proportion of events in New Mexico and Arizona (Figure 5B). With prior fuels management and indirect suppression tactics aimed at altering fire behavior, which can influence post-fire severity, these events represent potential opportunities to meaningfully increase the amount of good wildfire on the landscape: shifting fire characteristics such that they aligned with desired outcomes in an additional 25% of the footprint already burned by these fires would have added an additional 318,000 hectares of good wildfire or about 23% of the area treated with prescribed fire over our study decade, a relatively small, but ecologically meaningful, amount when compounded decade after decade.

**Figure 5.**
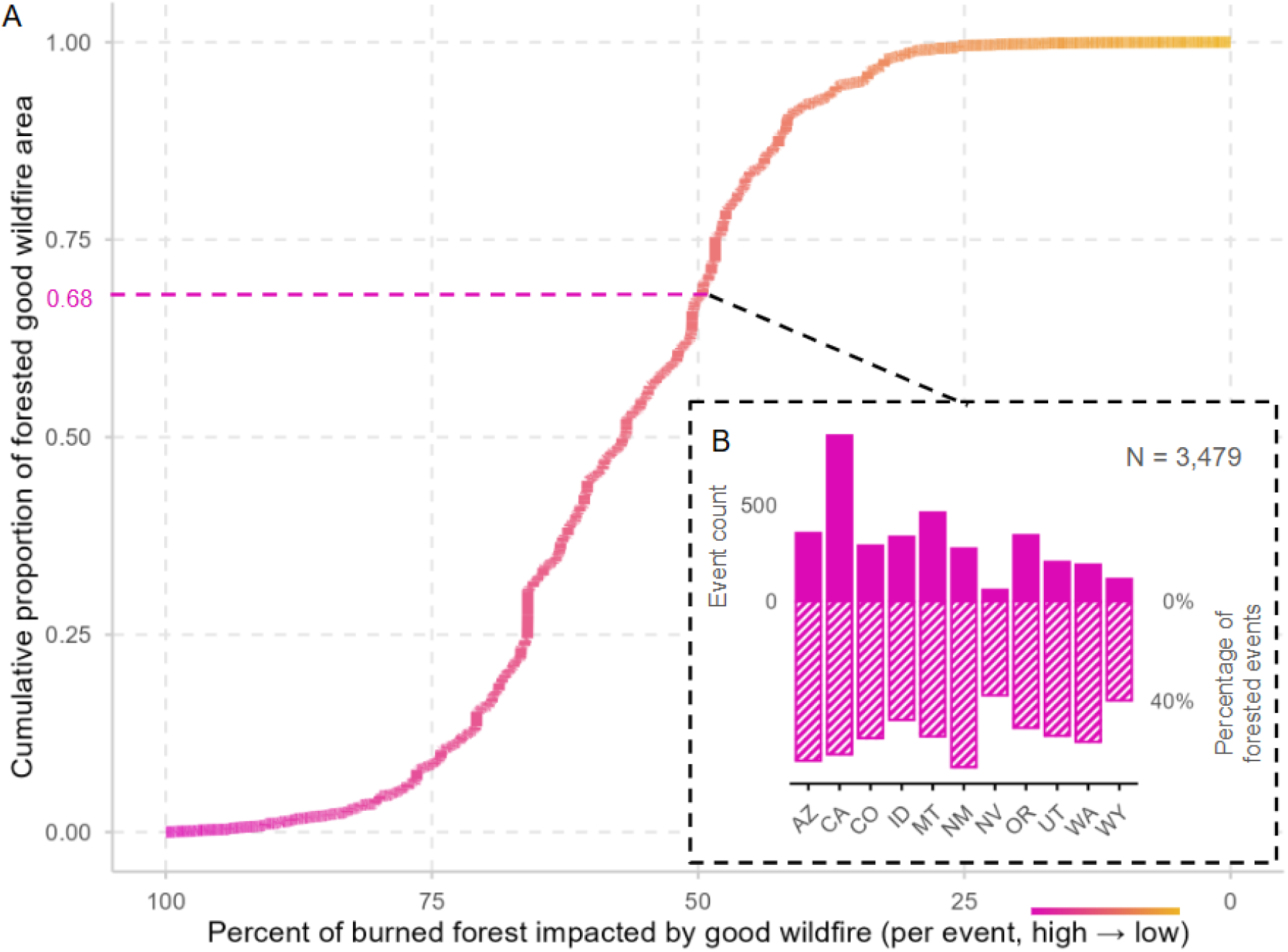
**(A)** Cumulative proportion of forested good wildfire area for events with >10% forest, by event percentage of good wildfire (note, x-axis descends from high to low).The pink to orange color scale represents the same x-axis in a more visual format. Pink dashed lines and accompanying text indicate the cumulative area proportions for events with greater than 50% good wildfire. **(B)** The number of events (top y-axis) and percentage of events (bottom y-axis) with greater than 50% good wildfire in each state (N = 3,479 total).

In order to achieve more area burned, as called for by recent federal and state land management policy (30, 31), we need to cultivate many forms of fire—from Indigenous cultural burning to prescribed fire to managing good wildfires. They all have potential ecological and social benefits and functionally help to build a mosaic of different fire severities and resulting forest densities and ages across a landscape. Indigenous scholars conceptualize “good fire” to refer to cultural burning and the traditional knowledge that encourages broader Indigenous fire stewardship (20, 32). Indigenous communities around the world have practiced forms of patch mosaic burning for millennia to reduce future fire size (33), mitigate the potential for extreme events [(34)], and manage ecosystems for cultural purposes (35, 36) and biodiversity (37). In the U.S., it is noteworthy that the Bureau of Indian Affairs was the only federal agency to substantially increase intentional burning between 1998 to 2018 (38).

At a national scale, there is policy and management imperative to fundamentally conduct more prescribed burning (38, 39). The intended $4.4B federal investment in the U.S. for fuel reduction treatments including prescribed fire by the Infrastructure Investment and Jobs Act (IIJA, 2021) and the Inflation Reduction Act (IRA, 2022) (40) was an important step to achieve these goals, though some of these funds were rescinded with the passage of recent federal legislation (41). Federal land managers are accomplishing significant amounts of prescribed burning (Figure 2) but need to treat much more area to meet restoration goals. One underrecognized way is through expansion of managed wildfire use (i.e., any strategy other than full suppression) (42). Land management agencies should get credit for the hard work that actually contributes to wildfires burning at lower severities. While we only consider fire severity and return interval here, there are many types of good wildfire that should be considered in future research, such as slower and cooler fires, which will be easier to manage. Fundamentally, we need to recognize that wildfires themselves are both reducing future fire risk and regulating ecosystem structure and composition at broad scales. To reduce barriers to more use of wildfire, we need to confront the fundamental assumptions of U.S. federal environmental policies that fail to recognize the importance of fire in fire adapted ecosystems and instead regard fire as a threat to resources (43).

While we explore two aspects of good wildfire, severity and fire return interval, there are many characteristics that are worth exploring in future work. There is a key need to develop a framework for measuring multiple dimensions of beneficial wildfire that enables managing for ecological and social benefits. For example, good wildfires may also be longer-interval, slower, smaller, patchy, and/or easier to manage during suppression. Good wildfires may have more specific ecological benefits, such as invasive species management (44), disrupting herbivory and predation of native species (45), or promoting re-sprouting of grasses for livestock or hunting (46). We may also define good wildfires with a combined social-ecological emphasis (47), integrating ecological benefits in tandem with minimized impacts on society and infrastructure, such as reducing risk (48), homes and lives lost, or smoke impacts (49).

Knowing that wildfires have always been inevitable in terrestrial ecosystems and will likely continue to increase across the western U.S., we need to fundamentally shift our management approach. We must cultivate good wildfires that will be more restorative, while at the same time protecting people and homes. There is the potential for fire management tactics to accomplish this, such as promoting lower intensities at the interior area of wildfires, rather than just focusing suppression on fire edges. We should capitalize on recent wildfire footprints as places to prioritize future prescribed and cultural burning and use wildfire boundaries as spatial breaks and control points for future burns that restore adjacent fire-excluded ecosystems (50). Capitalizing on wildfires in these and other ways offers promise for hastening the recovery of fire-prone landscapes across the western U.S. In a warming climate with more extreme wildfires projected (51), the window is shrinking for less intense and lower severity fires. We need to aim for a modern fire stewardship that accounts for good wildfire, implements substantial prescribed burns (38), and honors cultural burning practices (52), while still accommodating our settlement patterns within ecosystems that need fire. If we imagine that everywhere flammable in the western U.S. will burn—as a question of when, rather than if—how can we manage our landscapes? Seeing wildfires for the positive outcomes they can generate will allow land managers and society to better sustain fire-prone landscapes that are home to tens of millions of people.

## Data availability

All datasets used are publicly accessible. Fire event delineations were extracted from the combined wildland fire event dataset (54). Prescribed burn data was provided by the Treatment Wildfire Interagency Geodatabase (TWIG) (67). Raster datasets were accessed using Google Earth Engine, and include historic fire regime data from the LANDFIRE Fire Regime Groups (FRG)

(https://developers.google.com/earth-engine/datasets/catalog/LANDFIRE_Fire_FRG_v1_2_0) and Percent Replacement-Severity (PRS) datasets (v1.2.0)

(https://developers.google.com/earth-engine/datasets/catalog/LANDFIRE_Fire_PRS_v1_2_0), while remotely sensed data is from the Landsat series of satellites

(https://developers.google.com/earth-engine/datasets/catalog/landsat). Landcover data is from the Landscape Change Monitoring System (LCMS)

(https://developers.google.com/earth-engine/datasets/catalog/USFS_GTAC_LCMS_v2024-10) and Land Change Monitoring, Assessment, and Projection (LCMAP) data products

(https://gee-community-catalog.org/projects/lcmap/?h=lcmap#lcmap-level-1-land-cover-classes).GEDI footprint-based biomass density data were extracted from

https://developers.google.com/earth-engine/datasets/catalog/LARSE_GEDI_GEDI04_A_002.

## Code availability

All code used for analyses associated with this article are publicly available at: https://github.com/TylerLMcIntosh/a-number-on-good-fire.

## Methods

We analyzed wildfires from 2010-2020 within 11 states in the western United States: Washington, Oregon, California, Idaho, Montana, Nevada, Utah, Colorado, Wyoming, New Mexico, and Arizona. In order to capture wildfires of all sizes for our area and time frame of interest, we filtered the “combined wildland fire polygons” dataset from Welty and Jeffries (2020), which includes the Monitoring Trends in Burn Severity (MTBS) dataset (for fire perimeters ≥ 405 ha in the western U.S. starting in 1984) (53) in addition to data from 39 additional data sources, providing the best-available data on fires (< 404 ha) (54). This dataset is designed to be a comprehensive collection of wildfire boundaries within the United States. In total, our filtered dataset included a total of 18,061 wildfire perimeters for the western U.S.

Within fire perimeters and at prescribed burn event locations, we focused exclusively on areas that were forested one year before the fire. We used two land cover products to calculate forested area: the Landscape Change Monitoring System (LCMS) Land Cover Class v2022.8 (55) and the Land Change Monitoring, Assessment, and Projection (LCMAP) Primary Land Cover v1.3 (56). Areas were considered forested only if both land cover products classified them as “tree cover” (i.e., deciduous, evergreen, and mixed forest trees; woodlands were excluded) the year prior to the wildfire or prescribed fire event. Combining the two land cover maps produced a more conservative estimate of forest cover, and reduced the ‘false positive’ forest present in either individual product. Utilizing the combined dataset, the total forested area across the 11 western U.S. states was 62 million ha in 2020.

Within the forested pixels of each fire perimeter, we used Landsat imagery accessed on Google Earth Engine (57) to calculate an estimate of the Composite Burn Index (CBI), a commonly used and ecologically meaningful measure of fire severity. CBI was estimated using the method described in Parks et al. (2019), which uses random forest regression to calculate CBI based on Relativized Burn Ratio (RBR), latitude, climatic water deficit, and other factors. RBR was calculated using pre- and post-fire image composites of Landsat 4-9 imagery (Collection 2) during the growing season. A correction was applied to the CBI estimates to prevent overprediction at low values, resulting in a bias-corrected CBI (19). We applied commonly used CBI thresholds to identify low-moderate (CBI ≥ 0.1 and < 2.25) and high severity burned areas (CBI ≥ 2.25), e.g., see (58). High severity CBI is associated with near complete vegetation mortality, while low-moderate CBI is associated with lower levels of vegetation cover change and mortality (58). Fire severity estimates were computed at the annual temporal scale due to accepted practices of using year-prior to year-after imagery for remotely sensed burn severity calculations. As a consequence, our estimates treat areas burned twice in one year as burned in a single event, as there is no way to disentangle the fire severity effects of one fire versus a later fire in the same year. This avoids double-counting of area burned in regime-matching fire multiple times in one year, but will, by definition, also under-estimate the area burned at too high of a frequency for the regime. To validate that this decision has negligible impact on our results, we compared estimates of total forested area burned at the event level (multi-counting annual overlap) to estimates from the region level (annual overlap collapsed), and found that the estimates were within 0.004% of one another.

Based on a large body of literature (see main text, in addition to, e.g. (59–61)), we assume that contemporary wildfire in fire-adapted forested ecosystems will provide some level of ecological benefits when its severity level is aligned with the area’s historical fire severity and fire return interval (FRI) prior to European settlement. To estimate historical fire severity and frequency characteristics, we used the Landfire Historical Fire Regime suite of products (v1.2.0), which models presumed fire regime characteristics based on vegetation dynamics, fire behavior and effects, and biophysical context (62). The Landfire Fire Regime Groups (FRG) dataset (63) is derived by applying thresholds to the more granular Fire Return Interval (FRI) and Percent Fire Severity (PFS) datasets. All three datasets represent the cumulation of over 900 vegetation dynamic models and the input of over 800 experts, providing the best-available wall-to-wall estimates of historical fire regimes available for the United States (64). To identify areas with return intervals long enough that fires documented in the modern record would represent a frequency higher than historically expected, we applied the standard thresholds (i.e. used in FRG) of ≤35 years, >35 years, and >200 years to the FRI dataset (62). To distinguish between areas of historical “low and mixed severity” and “replacement severity” we applied the standard threshold (i.e., used in FRG) of 66% replacement severity fire to the PFS dataset, specifically the Percent of Replacement-Severity Fire (PRS) layer (62). The combination of these thresholds resulted in six fire regime classes, four of which are equivalent to FRG classes 1-4, and two of which represent FRG class 5 split into low and mixed severity (“FRG 5L”) and replacement severity (“FRG 5H”) (Supplementary Figure 1), as the standard FRG class 5 represents ‘any’ severity. We then identified two classes of good wildfire in forested ecosystems: low-moderate severity and high severity (Supplementary Figure 1). “Lower severity good wildfire,” (the primary focus of this research) was defined as low-moderate severity pixels within historically “Low and Mixed Severity” regimes. “High severity good wildfire” was defined as high severity pixels within historically “Replacement Severity” regimes. We then removed any identified good wildfire in regions with>35 year return interval and a documented prior wildfire that burned at an estimated bcCBI >0.1 at that location. Given these considerations, this approach resulted in a conservative estimate of wildfire occurrence between 2010 and 2020 with the potential for driving ecosystem benefits, as it does not fully account for the potential for mixed-severity fire driving benefits in mixed-severity regimes. Even in well-established regimes with more homogeneous severity, small patches of non-aligned severity wildfire may have ecological benefits such as increasing biodiversity or driving heterogeneity in forest structure and composition (65, 66). Here we do not address this dynamic, which is beyond the scope of this work. Our estimates are similarly conservative with regard to FRI. We do not account for the fact that fire regimes represent average conditions; shorter-than-average fire return intervals, or very low severity return in high severity regimes may in fact be within a historical range.

To compare estimates of good forest wildfire to estimates of forested area impacted by prescribed burning in the western U.S., we used data on 24,817 prescribed burn events in forest systems on national public lands from the Treatment Wildfire Interagency Geodatabase (TWIG) dataset (67). Prescribed burning of forested private lands, which is less common than prescribed burning on federal lands (68), is not accounted for. Further, prescribed burning on state lands is also not included. The TWIG dataset is currently the most comprehensive, accessible, and validated source of data on prescribed burning in the United States, incorporating data from the National Fire Plan Operations and Reporting System (NFPORS) (69) and Forest Activity Tracking System (FACTS) (70). The dataset was filtered to remove records flagged by the TWIG dataset creators as potential errors. For each documented treatment, we reduced the data to spatial centroids, as the use of polygons and calculated CBI would not be appropriate for analyzing prescribed fire given the diversity of forest treatments included in this analysis. We extracted LCMS, LCMAP, and Landfire Fire Regime Group data at each NFPORS data point in Google Earth Engine and processed the data under the assumption that the land cover and historic fire severity characteristics at the datapoint was representative of conditions across the entire burn area. Prescribed burns were only included in our analysis if treatments had an accomplishment date and both land cover datasets identified forested cover at their location in the year prior to treatment. We defined prescribed burning as inclusive of the following treatment types: broadcast burning, machine pile burning, jackpot burning, and hand pile burning. Fire use (including managed wildfire, backburning, and similar tactics) was excluded as the documented events had over 97% overlap with wildfire perimeters used for good wildfire calculations. Of all the forested prescribed burn events 22,057/24,817 (88.9%) were in low-moderate severity regions. The total area of all prescribed burns that was co-located with these same regions was 1,089,901 ha/1,406,828 ha (77.5%). When comparing the amount of good wildfire with the area of prescribed burns in low-moderate severity regions, wildfires burned 2.9 times as much area in low-moderate severity burning than the area impacted through prescribed burning.

To quantitatively describe recovery trajectories of burned areas, we used the aboveground biomass density (AGBD) metric derived from the Global Ecosystem Dynamics Investigation (GEDI) (71, 72). The AGBD metric for burned areas from all 18,061 fires were downloaded using GEDI Level 4A rasters in Google Earth Engine. GEDI Level 4A footprints were filtered using established quality criteria (l4_quality_flag = 1; degrade_flag = 0; AGBD > 0; sensitivity ≥ 0.90). Unburned control points were selected in locations within the same fire regime group that belonged to National Land Cover Database classes 41 (deciduous forest), 42 (evergreen forest), or 43 (mixed forest) for all years between 2008 and 2020, to ensure areas were forested prior to fire, that were beyond the fire boundary but within 500 m to 1000 m from the fire perimeter (73, 74). We excluded control points from the fire boundary into the first 500 m of unburned area to account for uncertainty in fire boundary delineations. We calculated summary statistics of mean, median, and standard deviation of AGBD per fire event, per severity class. We estimated percent biomass change with respect to the unburned biomass per fire event to evaluate the recovery trajectory based on the severity category of the fires within and beyond the fire regime group.

## Acknowledgments

We are grateful to participants at the Cary Institute and UCLA Townhall on Western Fire and Forest Resilience in September 2023 for discussions that informed this effort. This study was supported by: Earth Lab, University of Colorado - Boulder, Grand Challenge and CIRES (JKB, RVP); National Science Foundation Macrosystems Program grant 2017889 (JKB, TLM); National Science Foundation CAREER Program grant 1846384 (JKB, NTI, TLM); the Gordon and Betty Moore Foundation grant GBMF11974 and 13283 (WDH, JKB, CLR, BJH), Lyda Hill Philanthropies (WDH, JKB, CLR); and the Corbett Fire Endowment (JMV, KC). BJH acknowledges support from the Jack Corkery and George Corkery Jr. Endowed Professorship in Forest Sciences.

## Author Contributions

Conceptualization: JKB, WDH, KDM, RVP, CLR, BJH; Methodology: JKB, TLM, RVP; Investigation: JKB, TLM, RVP; Formal analysis: TLM, RVP; Visualization: JKB, TLM, NTI, MEB, RVP; Data curation: TLM, KC; Funding acquisition: JKB, WDH, BJH; Project administration: TLM, MEB, JKB; Resources: JKB; Supervision: JKB; Validation: TLM; Writing - original draft: JKB, TLM, RVP, KDM; Writing - review & editing: All authors

## Additional information

This article contains Supplementary Information. Correspondence and requests for materials should be addressed to Jennifer K. Balch.

## Supplementary figures

**Supplementary Figure 1.**
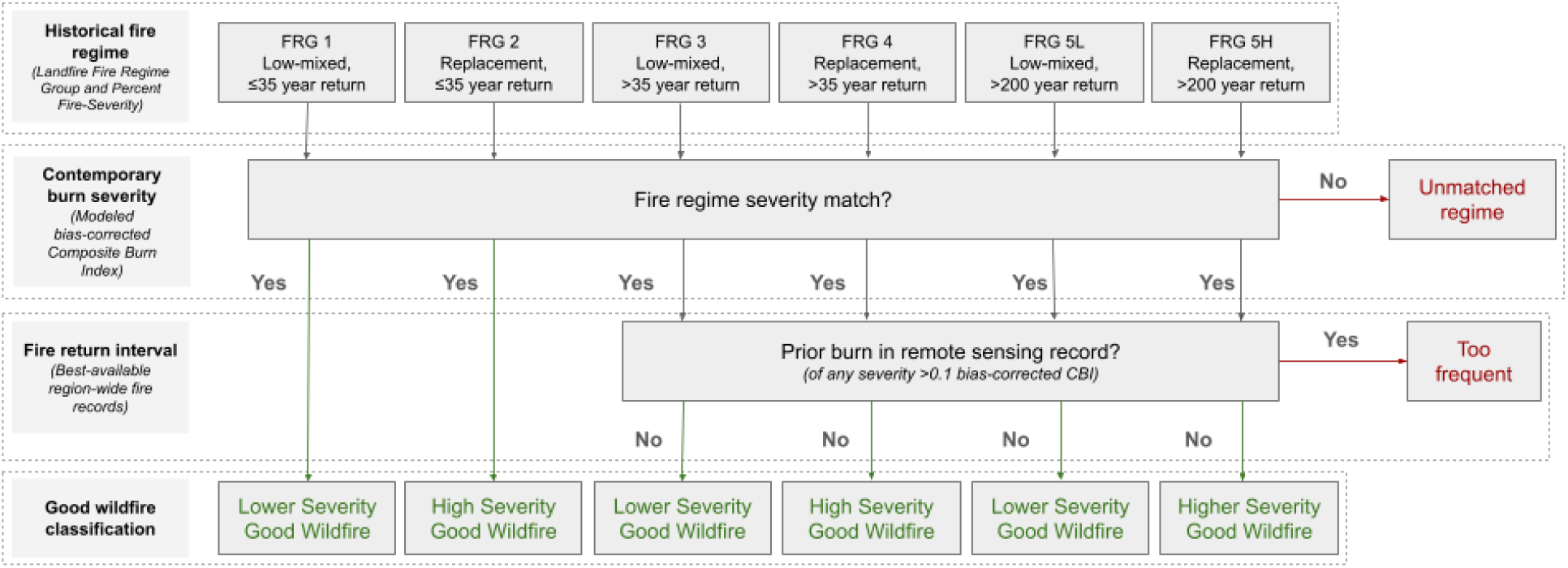
Good wildfire classification outline. Historical fire regime is established using the Landfire Fire Regime Group (FRG) and Percent Fire Severity (PRS) datasets to identify commonly used time frame and severity thresholds. Note that we split the Fire Regime class 5 into low-mixed (FRG 5L) and replacement (FRG 5H) regimes. Burn severity for contemporary wildfires is estimated using modeled bias-corrected Composite Burn Index (CBI) and compared to Landfire estimates of historical severity. For those fire regimes with >35 year return interval, the full remote sensing fire record is checked for additional wildfire at that location occurring with >0.1 bc-CBI.

